# Genomic Analysis of *Legionella pneumophila* in the Drinking Water System of a Large Building over 25 Years

**DOI:** 10.1101/2024.11.28.625380

**Authors:** Helena MB Seth-Smith, Adrian Egli, Sylvia Gautsch, Michael M. Bornstein, Eva M. Kulik

**Affiliations:** Institute of Medical Microbiology, University of Zürich, Zurich, Switzerland; Division of Clinical Bacteriology and Mycology, University Hospital Basel, Basel, Switzerland; Applied Microbiology Research, Department of Biomedicine, University of Basel, Basel, Switzerland; State Laboratory Basel-City, Basel, Switzerland; Department of Oral Health & Medicine, University Center for Dental Medicine UZB, University of Basel, Basel, Switzerland; Department Research, University Center for Dental Medicine UZB, University of Basel, Basel, Switzerland

**Keywords:** whole genome sequencing, dental institution, waterlines, evolution, ST45, water quality, surveillance, monitoring

## Abstract

*Legionella pneumophila,* the causative agent of Legionnaires’ disease, is often found in the plumbing systems of buildings from where it can be transmitted to humans via inhalation or aspiration of contaminated water drops. Annual routine water sampling from the potable water system of an occupational healthcare building in Basel over 25 years was performed in accordance with national guidelines. Overall, 309 water samples were collected at 38 time points over the period of 25 years. *L. pneumophila* was recovered from 120 water samples (38.8%) from 26 time points. No clinical infections were recorded during this period. Initial decontamination measures were successful for approximately 12 years, after which an increase in the total number of *Legionella* colony forming units as well as of *L. pneumophila*-positive sites was noticed, in 2008. Whole genome sequencing (WGS) analysis of n=123 isolates from n=113 samples showed all *L. pneumophila* to be sequence type (ST)-45 (Sequence based typing scheme). The isolates are closely related, with only 408 single nucleotide polymorphisms (SNPs) among all isolates after the bioinformatic removal of recombination events. Over the 25 years, a single lineage deriving from a recent common ancestor colonized the water system of this building. The phylogeny of isolate genomes can be interpreted as inferring good water circulation, possible recolonization from a common source after cleaning, with genome evolution and insertion / loss of large elements evident. Regular monitoring of waterlines in healthcare settings helps to identify concentrations of *Legionella* spp. and WGS is recommended for detailed investigation.

**Data Summary:** All data is submitted to the ENA under project PRJEB79004 under accession numbers ERR13662450-ERR13662572.

**Impact Statement:** This is the most detailed, long-term study of *L. pneumophila* in the water system of a single building recorded to date. The *L. pneumophila* isolates found in the building over the sampling period of 25 years were all closely related, belonging to ST45. SNP analysis suggested that the common ancestor of the cluster was from around 1938 (range 1911 to 1959), and movement of a large genomic island and plasmid transfer were observed. Despite several decontamination measures, it was impossible to completely eradicate *Legionella* spp. from the water system of the historic building. No infections could be attributed to the presence of *L. pneumophila* in this building. To mitigate the risk of Legionellosis from such buildings, awareness, regular water testing based on official national guidelines and recommendations, and other control measures, such as the use of sterile water for critical procedures can be recommended.

## Introduction

*Legionella pneumophila*, a Gram-negative, aerobic and non-spore-forming bacterium, is an opportunistic pathogen that can present a risk to human health. It can cause Legionnaires’ disease, a severe atypical pneumonia, or Pontiac fever, a milder form of the same infection (1). Of the over 60 known *Legionella* species, *L. pneumophila* serogroup 1 is responsible for most severe infections. Transmission from environment to human occurs through inhalation or aspiration of aerosols containing *L. pneumophila*, where the bacteria infect alveolar macrophages in the lung (2,3).

*L. pneumophila* are ubiquitous in various freshwater habitats as well as man-made water systems such as potable water systems, fountains, spa water, and air-conditioner cooling towers (1,4). In these environments, *L. pneumophila* can be found planktonically or coexisting with other microorganisms in biofilms where they can parasitize and replicate within free-living amoebae (5). The association with amoebae offers benefits to *Legionella*, providing protection against disinfectants, UV radiation, or fluctuations in water temperature (6). Water temperatures between 25°C and 45°C are optimum for growth and may reach critical concentrations. At higher water temperatures, bacterial growth is inhibited and temperatures above 60°C start to be bactericidal for *L. pneumophila*.

Due to this ability to grow within pipes, guidelines and regulations exist to control the growth of *Legionella* spp. in the water systems of buildings. In addition to sanitary measures such as regular maintenance of the waterlines or temperature checks, regular microbiological analysis of water samples is recommended to monitor the water quality (7,8). This also applies to healthcare facilities, such as hospitals or clinics which may treat vulnerable persons having a potential higher risk of acquiring Legionnaire’s disease. In dental institutions, biofilm formation in dental unit water lines may result in high numbers of microorganisms including *Legionella* spp. in the water used for cooling or ultrasonication procedures (9). Consequently, an increased risk for *Legionella* infection has also been assumed for dental health care workers; however, a recent review suggests that there is only very limited evidence for an increased risk in this professional group (10).

Epidemiological studies based on the analysis of genomic data are very helpful in identifying transmission routes and tracing potential sources of infections. In addition, such studies can also be useful to assess the microbial diversity and evolutionary relatedness in various environments. Genomic epidemiological analysis, together with aggressive interventional measures, were central to control an outbreak of nosocomial cases of *L. pneumonia* in an Australian hospital (11). In this case eradication methods, which included disinfection of the water distribution system with a chlorinated, alkaline detergent as well as removal of redundant plumbing system, appeared to be successful in eliminating the *L. pneumophila* population responsible for the infections in the hospital for at least six months. However, other studies show that artificial water systems remain colonized by *L. pneumophila* despite the implementation of decontamination measures such as the superheat and flush method (12,13).

At the Dental School of the University Basel, Switzerland, routine water testing began in the early 1990s with a focus on tap water, to assess and control bacterial contamination at the source level before it could enter the waterlines of dental units. In this context, a unique collection of *L. pneumophila* strains isolated from different points in the water system was created, before and after decontamination measures, in the same water distribution system, from 1994 to 2018. In summer 2019, the Dental School of the University Basel moved to a new building in a different location.

The aim of this study was to investigate the distribution and evolution of *L. pneumophila* within a single building, over 25 years.

## Methods

### Study setting and sample collection

Routine testing of the drinking water system was performed annually at the Dental School of of the University Basel, Switzerland at a regular basis. The building has housed the Dental School of the University Basel for almost one hundred years and has been, if necessary, modified to meet new requirements during this period. In particular, the historic part of the building contained older water pipes and was not initially designed to prevent potential colonization by *Legionella* spp.

Therefore, the sampling points were selected based on a risk assessment and included end points of the water distribution system as well as rarely used faucets. Additional tests were carried out as required and after consultation with the Hygiene Commission of the building. Water samples were usually taken from the warm water circuit and processed in accordance with national guidelines that were in force at the respective time (8). Following species identification, *L. pneumophila* isolates were stored at −70°C until analysis. As required by the guidelines, 200 ml of water was taken from the respective water outlet and filtered by pressure through a membrane filter. The filter was then placed onto an agar plate selective for *Legionella* species. When reporting the total colony-forming units (CFU), numbers are calculated to one liter of water.

### Culture and DNA extraction

*L. pneumophila* strains isolated during the routing water tests in the years 1994 to 2018, i.e. during a period of 25 years, were included in this study. The isolates were grown on Buffered Charcoal Yeast Extract (BCYE) Agar (Thermo Fisher Scientific, Reinach, Switzerland) aerobically in a humidified atmosphere at 37°C for 48-72 h. DNA was extracted using the Qiagen EZ1 with the DNA tissue kit (Qiagen AG, Hombrechtikon, Switzerland).

### Genome sequencing and analysis

All samples were sequenced on the NextSeq500 PE150 after Illumina DNA prep, resulting in over 30x mean read depth per sample. Isolate ZIB8567 was sequenced by Oxford Nanopore Technologies (ONT) on an R10.4 flowcell to a mean read depth of 50x and base called with Dorado hac@v4.2.0 (https://github.com/nanoporetech/dorado). All data is submitted to the ENA under project PRJEB79004.

Ridom v 9.0.10 was used on Unicycler v0.3.0b (10.1371/journal.pcbi.1005595) assemblies to assign Sequence Based Typing (SBT) sequence types (ST) to the isolates. Data on two ST45 samples (917837 and 919569) were obtained from the UK Health Security Agency (UKHSA) and used as outgroups.

For phylogenetic analysis, the hybrid Unicycler v0.4.8 assembled genome of isolate ZIB8567 (isolated in 2018), comprising a single contig of 3522222 bp was used as the reference against which to map the reads of all isolates within CLC Genomics Workbench v22.0.2, calling variants with parameters: variant calling with 10x minimum coverage, 10 minimum count and 70% minimum frequency. Consensus genomes were extracted from mapped reads using a minimum coverage of 10. A multiple sequence alignment (MSA) was created from the whole genome alignments, exported and run through Gubbins v3.3.0 (14) with five iterations to remove recombinations. Results were viewed in Phandango (15). BactDating v1.1.1 (16) was run four times on the Gubbins output with one million iterations and the arc model to check the reproducibility, and representative results are given. Abricate v1.0.1 (https://github.com/tseemann/abricate) and the NCBI database (17) were used to define resistance determinants present.

Regions of the genome present in the sequenced genomes and absent from the genomes of strain Philadelphia (SRR801743 and ERR351242), or present at very low coverage, were identified by analyzing the mapping statistics of all isolates in CLC (<99.9% of reference mapped to), visualizing mapped reads in Artemis, and marking regions without mapping. Regions over 1kb were compared with blastn against the nt database to check whether there were matches in other submitted genomes, and top matches are given. Insertion elements (ISs) were categorized using ISFinder (https://isfinder.biotoul.fr). Further elements were characterized by iterative analysis of the mapping statistics (<98% reads mapped) and *de novo* assembly of Illumina data in CLC using default parameters (slow) and mapping of reads as above.

## Results

### Sampling

Overall, 309 water samples, taken from different locations in the building, were collected at 38 time points over a period of 25 years. As the building, including waterlines, was renovated over the course of the 25 years at various locations, the same sampling points might not have been accessible any longer and therefore, nearby faucets fed by the same waterline were selected. *L. pneumophila* was recovered from 120 water samples (38.8%) and 113 *L. pneumophila* isolates from 26 time points could be included in this study (Figure 1), as well as 10 replicate isolates picked from the same plates as five of the isolates. After initial decontamination measures that were successful for approximately 12 years until 2008, assessed by presence and quantity of colony forming units (CFU), an increase in the total number of *Legionella* CFU as well as of *L. pneumophila*-positive sites was noticed (Figure 1).

**Figure 1.**
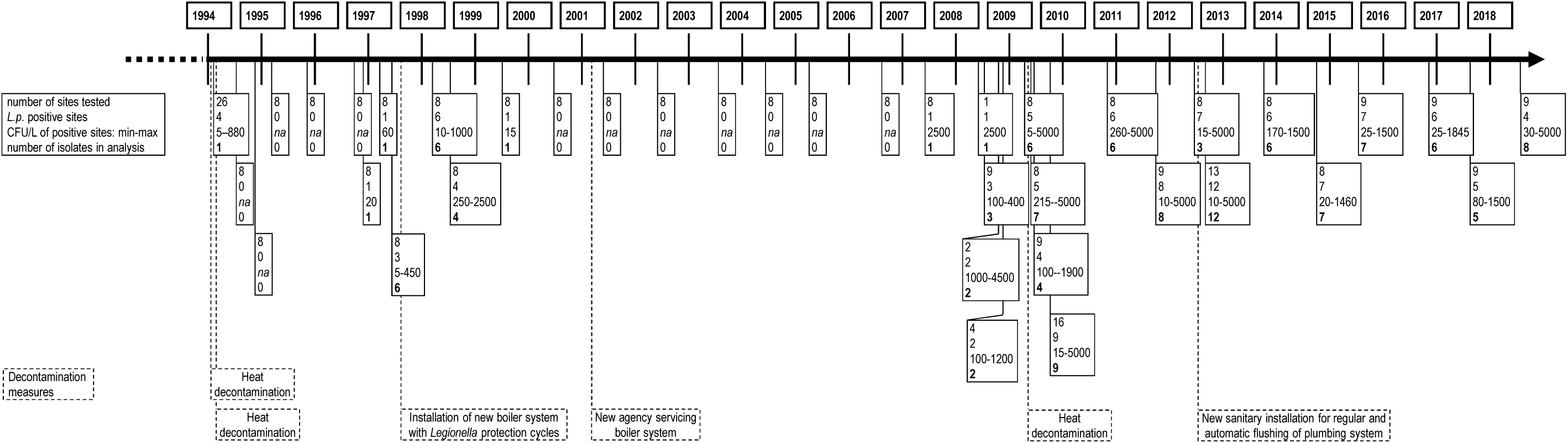
Timeline of *L. pneumophila* strains isolated over 25 years. Shown are the number of sites tested per time point, the number of sites that were *L. pneumophila* positive, the minimum and maximum CFU/L values for *L. pneumophila*-positive site per time point and the number of isolates per time point that was analysed (in bold) as well as the decontamination measures carried out during this time-period.

### Phylogenetic analysis of ST45 cluster

All 123 isolates from the building belong to ST45, which is within serogroup 1. They are unrelated to previously published samples from other locations within the City of Basel, where the Dental School of the University Basel is located (18).

Phylogenetic analysis of the 123 isolates, based on a complete genome hybrid assembly reference from an isolate within the cluster, shows that they are closely related (Figure 2). We found 408 SNPs across the whole cluster, with recombinations removed. There is no apparent clustering of isolates taken from the same sampling points, neither is there clear clustering between the replicate colonies taken from the same samples. A mutation rate of 0.38 SNPs per genome per year [range 0.27-0.49] and a most likely date for the common ancestor of this cluster of 1938 [range 1911-1959] was calculated using BactDating. The two ST45 isolates submitted to the UKHSA originating from Oxford in 2009 (917837) and of unknown provenance (919569) are respectively 1875 and 1888 SNPs from the ST45 isolates sequenced for this study (recombinations removed). The recombinations identified within the cluster represent 94% of the identified SNPs (6876/7284), although many fall within repeat genes and may represent mapping difficulties (Figure S1).

**Figure 2.**
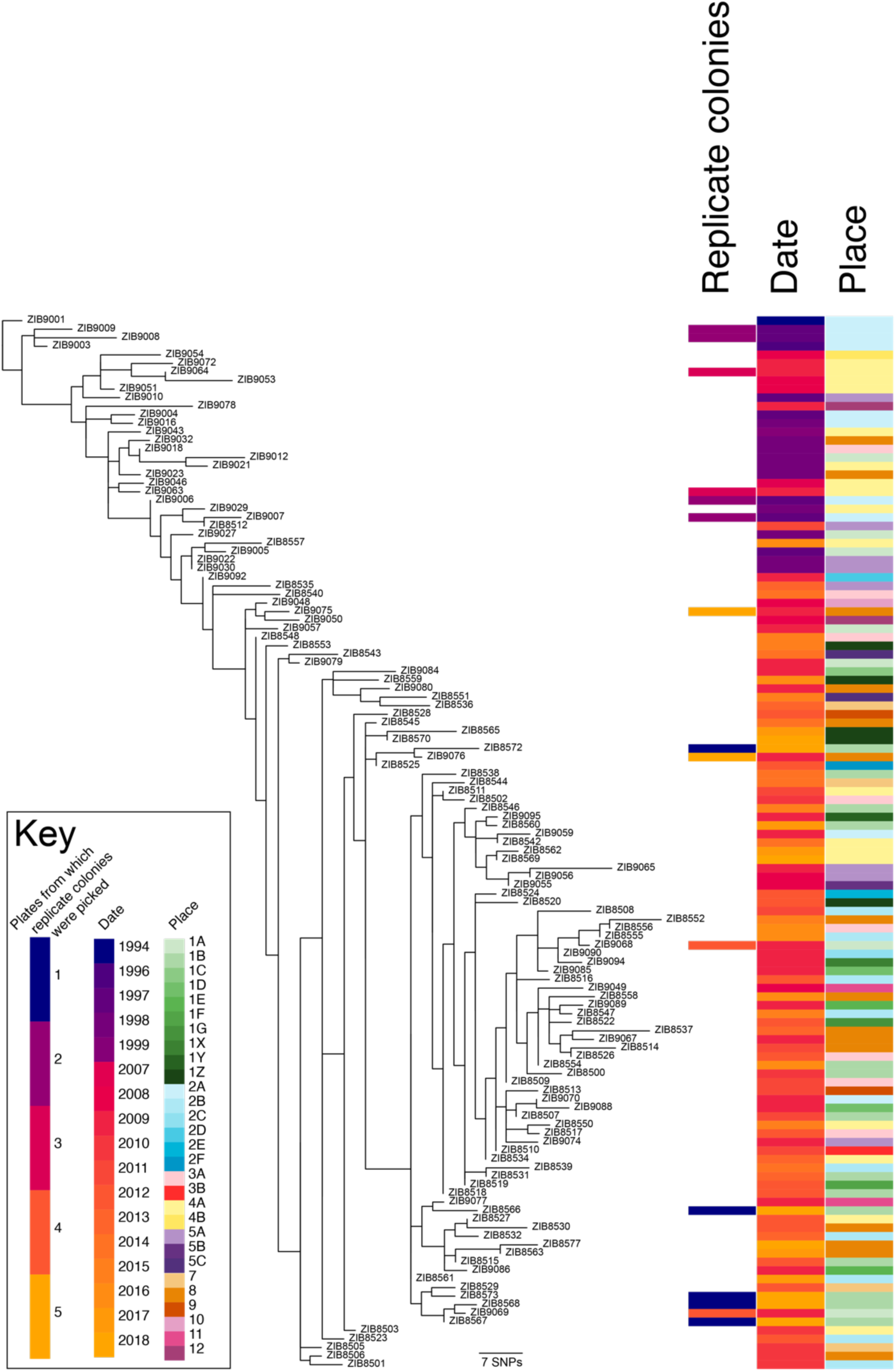
Recombination-adjusted phylogenetic tree of all isolates with associated metadata. Gubbins generated phylogeny based on mapping of all data against a complete hybrid assembly of ZIB8567, rooted on the oldest isolate (ZIB9001, 1994). Metadata (right) shows (left to right) which replicate isolates came from the same plates (replicate colonies), year of isolation (date) and location of isolation within the building (place). As some of the same sampling points were not always accessible during the 25 years, nearby faucets fed by the same waterline were selected and/or additional water outlets were included. Water outlets fed by the same waterline are represented by the same number. Figure generated using Phandango (15).

Regarding resistance determinants, the gene aph(9)-Ia, a chromosomally-encoded aminoglycoside phosphotransferase, was identified in the hybrid assembled genome of *L. pneumophila* ZIB8567 with a nucleotide identity of 89.81%.

**Figure S1.**
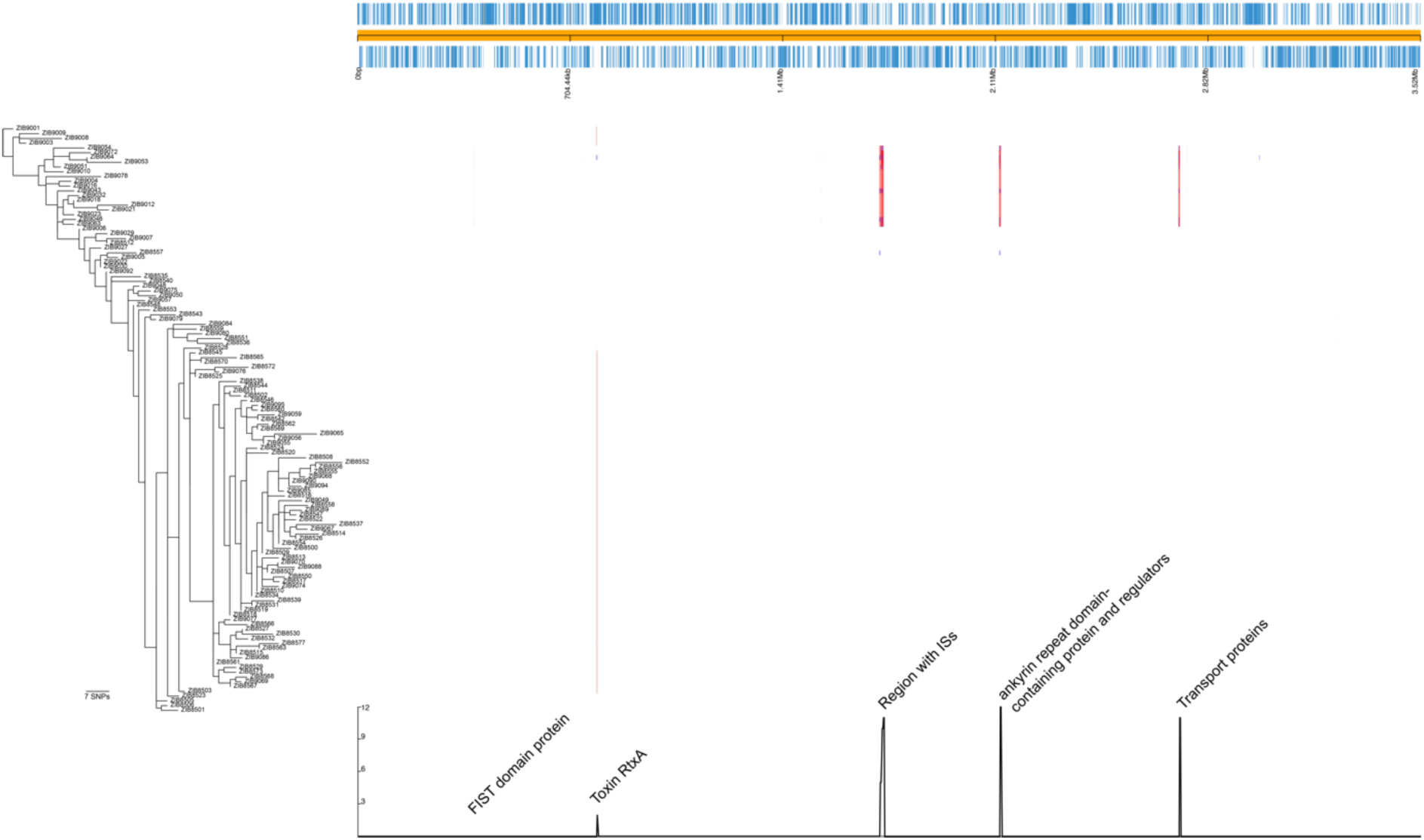
Recombinations identified within the ZIB cluster. Gubbins identified recombinations are shown aligned with the phylogeny rooted as in Figure 2. Genome annotation above the recombinations shows the coding sequences (CDSs) in blue along the chromosome. Below the recombination tracks, where recombination loci in single isolates are shown in blue, and those identified in more than one isolate are shown in red, is a graph of frequency with key locations annotated. Figure generated using Phandango (15).

### Regions of difference and genomic islands

All the presented *L. pneumophila* genomes share very similar gene content, with reads from all genomes mapping to over 99% of the ZIB8567 hybrid assembly. This is also the case for the ST45 genomes 917837 and 919569 obtained from UKHSA. Reads from *L. pneumophila* strain Philadelphia, however, cover only 87% of the assembly, with under 90% of the reads mapping.

Many regions of difference (RoD) were identified in the ST45 genomes which are not present in the Philadelphia reference strain genome (Table). When comparing to the two ST45 database genomes, only one possible genomic island (GI) appears to the unique to the isolates from this study: a 30kb region (172917-203527 in ZIB8567 hybrid assembly) of high read depth, sharing 90.55% identity over 92% of its length with a region of the genome of *Legionella longbeachae* strain B1445CHC (CP045306.1). This RoD carries phage-like genes with a Type IV secretion system cargo. This RoD is absent from the genomes of six isolates, namely ZIB8547, ZIB9003, ZIB9004, ZIB9016, ZIB9008, and ZIB9009, all originating from close locations 2A and 2B. The position of these isolates in the phylogeny infers that the ancestral strain of this cluster carried the RoD, and it had been lost on three independent occasions within the phylogeny.

All the rest of the RoD between ST45 and Philadelphia are present in the genomes of other strains of *L. pneumophila*, with most matches to that of strain Lorraine (FQ958210.1). Forty of the RoDs appear to be IS elements, of which the vast majority (n=38) are related to ISLpn9 (Table).

The isolates ZIB8557, ZIB9043, ZIB9046, ZIB9051, ZIB9053, ZIB9054, ZIB9063, ZIB9064, and ZIB9072, all from locations 4A and 4B, carry a 150kb plasmid identical to pB3526CGC_150k from *L. longbeachae* (CP042253.1)(19). According to the phylogeny, this suggests acquisition of the plasmid on three separate occasions.

## Discussion

*L. pneumophila* has been reported from the water systems of large buildings and hospitals in studies from Canada (20), USA (12,21,22), Italy (23), Germany (24), Finland (13), Poland (25), Scotland (26), and Australia (11). Genome sequencing has previously been used in such studies (11,27,28), but none over such a long period of time as in our study.

Our data shows a long-term presence of *L. pneumophila* ST45 within an old building in Basel in which the Dental School of the University Basel was located from 1924 to 2019. The estimated time of the most recent common ancestor of the cluster, at around 1938, would therefore fit with the age and development of the building as this historical building has been regularly renovated and new annexes added (29). While some buildings have been found to contain multiple STs (28), our study describes the same ST found in all locations at all timepoints, suggesting ongoing colonization of the waterlines with the same clonal strain over 25 years. That there was no clear phylogenetic signal related to location within the building, infers circulation through the waterlines, and less spatial structuring than described by David et al. (28). Likewise, no influence of the decontamination measures could be noted in the phylogeny in terms of clear termination of certain lineages. It is possible that a more in-depth analysis including more isolates from each timepoint might have resulted in a more nuanced picture in terms of clonal extinction, as decontamination measures have been shown to reduce the number of *Legionella* present (30,31). This might also have helped us to capture the presence of any *L. longbeachae* isolates.

Investigation of global data provided by UKHSA and from the literature shows that *L. pneumophila* ST45 has previously been isolated in environmental samples from China, Japan, South Korea, France, Germany, Belgium, and Italy, and in clinical isolates (n=41) from the USA, Canada, and across Europe including from two locations in Switzerland and the neighbouring state in Germany (Baden-Württemberg) between 2010-2013 (32–37). Very few genomes are available for comparison, and the two ST45 genomes available were over 1800 SNPs away from the presented cluster. Isolates within this ST have been found to cause clinical disease, yet no infections were identified within our study. It is commonly assumed that dental personnel have an elevated occupational risk for *Legionella* infection as they might potentially inhale contaminated aerosols generated by water-cooled rotary instruments used in dentistry. Based on antibody levels, it was suggested in 1985 that the risk of *Legionella* infection for dental personnel increases proportionally with increased clinic exposure time (38). However, no cases of *Legionella* infection were reported over the course of our study period among dental personnel with direct patient contact working at the Dental School of the University Basel (data not shown). This is in accordance with a recent meta-analysis that concluded that the occupational risk for an occupationally acquired *Legionella* infection for dental personnel is lower than previously anticipated (10). A difference was noticed between studies published prior to 1998 and those after 1998, attributed to the regular release of infection control guidelines in dentistry since being initiated at the turn of the century (10). Strict compliance with currently valid guidelines may have contributed to the absence of *Legionella* infections among dental personnel at the Dental School of the University Basel. Similarly, while contaminated water might be a potential risk for immunocompromised patients who encounter the aerosol, sterile water was generally used for this group of patients and the water from dental units was analyzed on a risk basis. In these tests, no *Legionella* spp. were detected except for *L. geestiana*, which was identified on one occasion in a dental unit in 2009 (data not shown). Nevertheless, awareness, regular testing, and other control measures, e.g. the use of sterile water for critical procedures are recommended.

The calculated mutation rate within this cohort of 0.38 SNPs per genome per year compares well to previous estimates of 0.71 SNPs per genome per year (David 2016) and 0.49 SNPs per genome per year (Sanchez-Buso 2014). *L. pneumophila* is known to be a recombinogenic species (39,40); the present study shows limited recombination, as is likely in an environment with limited genomic diversity, analysis of which may be confounded by repetitive genes. Recombinations around *hemA*, identified as hotspot in ST1 (40), were not identified in this ST45 cluster. Plasmid movement and genomic island loss were also observed, showing that genome plasticity is possible in this population. Interestingly, these events are associated with isolates from particular locations; however, it is not possible to infer specific selective pressure on these elements within these niches. Genetic elements of *L. longbeachae* could be detected, although this bacterium itself was not isolated in this study. Other *Legionella* species, such as *L. geestiana* and *L. anisa*, were detected albeit rarely (data not shown).

In conclusion, the present work demonstrates that a single clonal lineage of *L. pneumophila* can, despite decontamination measures, persist in a building’s water system for over 25 years. The phylogeny of the isolates can be interpreted as inferring good water circulation, and possible recolonization from a common source after cleaning.

## Ethical approval and consent to participate

No clinical data is described, and all isolates are environmental. As such, no ethical approval is required.

## Author contributions

HSS: Methodology, Software, Validation, Formal analysis, Data Curation, Writing - Original Draft, Visualization

AE: Resources, Supervision, Writing - Review & Editing, Project administration

SG: Resources, Writing - Review & Editing

MMB: Resources, Supervision, Writing - Review & Editing, Project administration

EMK: Conceptualization, Methodology, Validation, Formal analysis, Investigation, Resources, Data Curation, Writing - Original Draft, Visualization, Supervision, Project administration

## Acknowledgements

The authors are grateful for the excellent technical assistance by Krystyna Lenkeit and Irene Schweizer and for the valuable support of the former heads of the Hygiene Commission, Jürg Meyer and Tuomas Waltimo. Many thanks to the UKHSA, UK for supplying data on ST45 samples submitted to them. Many thanks to the technicians at University Hospital Basel for excellent technical assistance with the sequencing.

## List of Tables

**Table.** List of genomic islands identified in the isolates.

**Table. List of genomic islands identified in the isolates**
| Start | End | Length | Found in | Blastn query cover | Blastn Percent Identity | Match to Strain Accession | Comments |
| --- | --- | --- | --- | --- | --- | --- | --- |
| 57502 | 60351 | 2850 | All ST45 genomes | 100.00 | 99.79 | Lorraine FQ958210.1 | Homology to ISLpn9 |
| 65009 | 66795 | 1787 | All ST45 genomes | 100.00 | 100.00 | Lorraine FQ958210.1 |  |
| 69573 | 75226 | 5654 | All ST45 genomes | 100.00 | 100.00 | Lorraine FQ958210.1 |  |
| 107571 | 108853 | 1283 | All ST45 genomes | 100.00 | 100.00 | Lorraine FQ958210.1 |  |
| 154181 | 155180 | 1000 | All ST45 genomes | 100.00 | 99.60 | Lorraine FQ958210.1 | Homology to ISLpn9 |
| 172917 | 184420 | 11504 | Unique to ZIB genomes | 80.00 | 89.22 | Legionella longbeachae strain B1445CHC CP045306.1 | present at much higher coverage |
| 197488 | 203527 | 6040 | Unique to ZIB genomes | 100.00 | 98.24 | Legionella longbeachae strain B1445CHC CP045306.1 | present at much higher coverage |
| 203528 | 228760 | 25233 | All ST45 genomes | 100.00 | 100.00 | Lorraine FQ958210.1 |  |
| 254469 | 255506 | 1038 | All ST45 genomes | 100.00 | 100.00 | Lorraine FQ958210.1 |  |
| 258026 | 259597 | 1572 | All ST45 genomes | 100.00 | 100.00 | Lorraine FQ958210.1 |  |
| 261477 | 263700 | 2224 | All ST45 genomes | 100.00 | 99.69 | Lorraine FQ958210.1 | Homology to ISLpn9 |
| 310472 | 311473 | 1002 | Very low coverage in Phildelphia reference | 100.00 | 99.80 | Lorraine FQ958210.1 | Homology to ISLpn9 |
| 333355 | 336147 | 2793 | All ST45 genomes | 100.00 | 99.96 | Lorraine Q958210.1 |  |
| 359496 | 360497 | 1002 | All ST45 genomes | 100.00 | 99.90 | Lorraine FQ958210.1 | Homology to ISLpn9 |
| 364046 | 366198 | 2153 | All ST45 genomes | 100.00 | 100.00 | Lorraine FQ958210.1 |  |
| 388083 | 390250 | 2168 | All ST45 genomes | 100.00 | 100.00 | Lorraine FQ958210.1 |  |
| 392528 | 393528 | 1001 | All ST45 genomes | 100.00 | 99.80 | Lorraine FQ958210.1 | Homology to ISLpn9 |
| 470948 | 471949 | 1002 | Very low coverage in Phildelphia reference | 100.00 | 99.90 | Lorraine FQ958210.1 | Homology to ISLpn9 |
| 492613 | 493614 | 1002 | All ST45 genomes | 100.00 | 99.80 | Lorraine FQ958210.1 | Homology to ISLpn9 |
| 523278 | 524733 | 1456 | Very low coverage in Phildelphia reference | 100.00 | 99.93 | Lorraine Q958210.1 | Homology to ISLpn9 |
| 578328 | 579663 | 1336 | All ST45 genomes | 100.00 | 100.00 | NCTC11985 LT906452.1 |  |
| 620762 | 622020 | 1259 | All ST45 genomes | 100.00 | 100.00 | NCTC11985 LT906452.1 |  |
| 622674 | 660984 | 38311 | All ST45 genomes | 75.00 | 98.01 | Corby CP000675.2 |  |
| 698965 | 699965 | 1001 | All ST45 genomes | 100.00 | 99.80 | NCTC11985 LT906452.1 | Homology to ISLpn9 |
| 743322 | 746237 | 2916 | All ST45 genomes | 100.00 | 100.00 | NCTC11985 LT906452.1 |  |
| 782582 | 791077 | 8496 | All ST45 genomes | 100.00 | 99.80 | NCTC11985 LT906452.1 | Across toxin gene *rtxA* |
| 795792 | 798713 | 2922 | All ST45 genomes | 100.00 | 100.00 | NCTC11985 LT906452.1 |  |
| 812799 | 818434 | 5636 | All ST45 genomes | 100.00 | 100.00 | NCTC11985 LT906452.1 |  |
| 925770 | 927658 | 1889 | All ST45 genomes | 99.00 | 97.78 | ERS1305867 CP048618.1 |  |
| 944897 | 946813 | 1917 | All ST45 genomes | 100.00 | 98.44 | Lorraine FQ958210.1 |  |
| 104851<br>2 | 105021<br>6 | 1705 | All ST45<br>genomes | 100.00 | 97.65 | ERS1305867<br>CP048618.1 |  |
| 105914<br>9 | 106077<br>6 | 1628 | Very low<br>coverage in<br>Phildelphia<br>reference | 100.00 | 99.63 | PATHC002<br>CP113439.1 | Homology to<br>ISLpn9 |
| 116532<br>6 | 116695<br>2 | 1627 | Very low<br>coverage in<br>Phildelphia<br>reference | 100.00 | 99.82 | Lorraine FQ958210.1 | Homology to<br>ISLpn9 |
| 118121<br>1 | 118298<br>2 | 1772 | All ST45<br>genomes | 100.00 | 100.00 | Lorraine FQ958210.1 |  |
| 119391<br>1 | 119550<br>3 | 1593 | All ST45<br>genomes | 100.00 | 100.00 | Lorraine FQ958210.1 |  |
| 120286<br>3 | 120538<br>0 | 2518 | All ST45<br>genomes | 100.00 | 99.80 | Lorraine FQ958210.1 | Homology to<br>ISLpn9 |
| 122745<br>1 | 123162<br>9 | 4179 | All ST45<br>genomes | 100.00 | 99.88 | Lorraine FQ958210.1 |  |
| 123335<br>8 | 123584<br>3 | 2486 | All ST45<br>genomes | 100.00 | 100.00 | Lorraine FQ958210.1 |  |
| 123770<br>8 | 123914<br>3 | 1436 | All ST45<br>genomes | 100.00 | 100.00 | Lorraine FQ958210.1 |  |
| 133302<br>3 | 133754<br>3 | 4521 | All ST45<br>genomes | 100.00 | 99.93 | Lorraine FQ958210.1 |  |
| 135544<br>3 | 135668<br>0 | 1238 | All ST45<br>genomes | 100.00 | 100.00 | Lorraine FQ958210.1 |  |
| 135859<br>9 | 136022<br>8 | 1630 | Very low<br>coverage in<br>Phildelphia<br>reference | 100.00 | 99.94 | Lorraine FQ958210.1 | Homology to<br>ISLpn9 |
| 143979<br>8 | 144109<br>3 | 1296 | Very low<br>coverage in<br>Phildelphia<br>reference | 100.00 | 99.46 | PATHC002<br>CP113439.1 | Homology to<br>ISLpn9 |
| 147732<br>7 | 147972<br>6 | 2400 | All ST45<br>genomes | 100.00 | 99.62 | PATHC002 CP113439 | Homology to<br>ISLpn9 |
| 159311<br>1 | 159442<br>8 | 1318 | Very low<br>coverage in<br>Phildelphia<br>reference | 100.00 | 99.85 | Lorraine FQ958210.1 | Homology to<br>ISLpn9 |
| 159560<br>9 | 159726<br>1 | 1653 | All ST45<br>genomes | 100.00 | 100.00 | Lorraine FQ958210.1 | Lower<br>homology to<br>ISLpn9 |
| 160594<br>2 | 160694<br>0 | 999 | All ST45<br>genomes | 100.00 | 100.00 | Lorraine FQ958210.1 | Homology to<br>ISLpn9 |
| 162725<br>4 | 162918<br>8 | 1935 | All ST45<br>genomes | 100.00 | 100.00 | NCTC11985<br>LT906452.1 |  |
| 167126<br>8 | 167743<br>0 | 6163 | All ST45<br>genomes | 100.00 | 100.00 | Lorraine FQ958210.1 |  |
| 172963<br>6 | 173275<br>0 | 3115 | All ST45<br>genomes | 100.00 | 100.00 | Lorraine FQ958210.1 |  |
| 174019<br>9 | 174129<br>9 | 1101 | All ST45<br>genomes | 100.00 | 99.64 | NCTC11985<br>LT906452.1 | Homology to<br>ISLpn9 |
| 175036<br>7 | 175200<br>8 | 1642 | Very low<br>coverage in<br>Phildelphia reference | 100.00 | 99.70 | Lorraine FQ958210.1 | Homology to<br>ISLpn9 |
| 177868<br>8 | 178102<br>9 | 2342 | All ST45<br>genomes | 100.00 | 100.00 | Lorraine FQ958210.1 |  |
| 186924<br>5 | 187127<br>5 | 2031 | All ST45<br>genomes | 100.00 | 100.00 | Lorraine FQ958210.1 |  |
| 189680<br>4 | 190087<br>6 | 4073 | Very low<br>coverage in<br>Phildelphia<br>reference | 100.00 | 99.80 | Lorraine FQ958210.1 | Homology to<br>ISLpn9 |
| 197181<br>8 | 197286<br>5 | 1048 | Very low<br>coverage in<br>Phildelphia<br>reference | 100.00 | 99.90 | Lorraine FQ958210.1 | Homology to<br>ISLpn9 |
| 201769<br>1 | 201895<br>3 | 1263 | All ST45<br>genomes | 100.00 | 100.00 | Lorraine FQ958210.1 | Homology to<br>ISLdr1 |
| 206102<br>1 | 206267<br>0 | 1650 | All ST45<br>genomes | 100.00 | 100.00 | Lorraine FQ958210.1 |  |
| 206722<br>6 | 207097<br>2 | 3747 | Very low<br>coverage in<br>Phildelphia<br>reference | 100.00 | 100.00 | Lorraine FQ958210.1 | Homology to<br>ISLpn9 |
| 208018<br>3 | 208196<br>2 | 1780 | All ST45<br>genomes | 100.00 | 100.00 | Lorraine FQ958210.1 |  |
| 210025<br>5 | 210248<br>0 | 2226 | Very low<br>coverage in<br>Phildelphia<br>reference | 100.00 | 99.91 | Lorraine FQ958210.1 | Homology to<br>ISLpn9 |
| 212722<br>4 | 213320<br>0 | 5977 | All ST45<br>genomes | 100.00 | 100.00 | Lorraine FQ958210.1 |  |
| 214741<br>3 | 215062<br>9 | 3217 | All ST45<br>genomes | 100.00 | 100.00 | Lorraine FQ958210.1 |  |
| 215216<br>2 | 215352<br>0 | 1359 | All ST45<br>genomes | 100.00 | 100.00 | Lorraine FQ958210.1 |  |
| 217824<br>5 | 218016<br>3 | 1919 | Very low<br>coverage in<br>Phildelphia<br>reference | 100.00 | 99.79 | Lorraine FQ958210.1 | Homology to<br>ISLpn9 |
| 219082<br>5 | 219768<br>1 | 6857 | All ST45<br>genomes | 100.00 | 99.97 | PATHC002<br>CP113439.1 |  |
| 221689<br>4 | 221789<br>5 | 1002 | Very low<br>coverage in<br>Phildelphia<br>reference | 100.00 | 99.90 | PATHC002<br>CP113439.1 | Homology to<br>ISLpn9 |
| 226613<br>9 | 226768<br>7 | 1549 | All ST45<br>genomes | 100.00 | 100.00 | Lorraine FQ958210.1 |  |
| 228606<br>0 | 228939<br>1 | 3332 | All ST45<br>genomes | 100.00 | 100.00 | Lorraine FQ958210.1 |  |
| 229271<br>0 | 231465<br>9 | 21950 | All ST45<br>genomes | 100.00 | 100.00 | Lorraine FQ958210.1 |  |
| 231500<br>9 | 231666<br>7 | 1659 | Very low<br>coverage in<br>Phildelphia<br>reference | 100.00 | 100.00 | Lorraine FQ958210.1 |  |
| 231911<br>7 | 232437<br>3 | 5257 | All ST45<br>genomes | 100.00 | 1005.00 |  |  |
| 2327309 | 2331362 | 4054 | All ST45 genomes | 100.00 | 100.00 | Lorraine FQ958210.1 |  |
| 2367178 | 2368459 | 1282 | All ST45 genomes | 100.00 | 100.00 | Lorraine FQ958210.1 |  |
| 2374126 | 2376841 | 2716 | All ST45 genomes | 100.00 | 100.00 | Lorraine FQ958210.1 |  |
| 2395108 | 2396866 | 1759 | All ST45 genomes | 100.00 | 100.00 | Lorraine FQ958210.1 |  |
| 2398121 | 2399129 | 1009 | Very low coverage in Phildelphia reference | 100.00 | 99.70 | Lorraine FQ958210.1 | Homology to ISLpn9 |
| 2437537 | 2438802 | 1266 | All ST45 genomes | 100.00 | 100.00 | Lorraine FQ958210.1 |  |
| 2530688 | 2531689 | 1002 | Very low coverage in Phildelphia reference | 100.00 | 99.80 | Lorraine FQ958210.1 | Homology to ISLpn9 |
| 2562295 | 2563296 | 1002 | All ST45 genomes | 100.00 | 100.00 | NCTC11985 LT906452.1 | Homology to ISLpn9 |
| 2624142 | 2697268 | 73127 | All ST45 genomes | 100.00 | 99.99 | Lorraine FQ958210.1 |  |
| 2700063 | 2708857 | 8795 | All ST45 genomes | 100.00 | 99.73 | Lorraine FQ958210.1 |  |
| 2711396 | 2713990 | 2595 | All ST45 genomes | 100.00 | 100.00 | Lorraine FQ958210.1 |  |
| 2720705 | 2728751 | 8047 | All ST45 genomes | 100.00 | 100.00 | Lorraine FQ958210.1 |  |
| 2755536 | 2756800 | 1265 | All ST45 genomes | 100.00 | 100.00 | Lorraine FQ958210.1 |  |
| 2760534 | 2762270 | 1737 | All ST45 genomes | 100.00 | 100.00 | Lorraine FQ958210.1 |  |
| 2774564 | 2776814 | 2251 | All ST45 genomes | 100.00 | 100.00 | Lorraine FQ958210.1 |  |
| 2784580 | 2788461 | 3882 | All ST45 genomes | 100.00 | 100.00 | Lorraine FQ958210.1 |  |
| 2871046 | 2873896 | 2851 | Very low coverage in Phildelphia reference | 100.00 | 100.00 | PATHC002 CP113439.1 | Homology to ISLpn9 |
| 2877959 | 2880331 | 2373 | All ST45 genomes | 100.00 | 99.96 | Lorraine FQ958210.1 |  |
| 2901276 | 2903059 | 1784 | Very low coverage in Phildelphia reference | 100.00 | 99.94 | Lorraine FQ958210.1 | Homology to ISLpn9 |
| 2927856 | 2930694 | 2839 | All ST45 genomes | 100.00 | 100.00 | Lorraine FQ958210.1 |  |
| 2938791 | 2941692 | 2902 | All ST45 genomes | 100.00 | 99.97 | Lorraine FQ958210.1 | Homology to ISLpn11 |
| 2944637 | 2949035 | 4399 | All ST45 genomes | 100.00 | 100.00 | Lorraine FQ958210.1 |  |
| 2949872 | 2951925 | 2054 | All ST45 genomes | 100.00 | 100.00 | Lorraine FQ958210.1 |  |
| 2967128 | 2974009 | 6882 | All ST45 genomes | 100.00 | 100.00 | Lorraine FQ958210.1 |  |
| 298089<br>6 | 298600<br>6 | 5111 | All ST45<br>genomes | 100.00 | 100.00 | Lorraine FQ958210.1 |  |
| 298883<br>8 | 299393<br>6 | 5099 | All ST45<br>genomes | 100.00 | 100.00 | Lorraine FQ958210.1 |  |
| 299666<br>0 | 300026<br>0 | 3601 | All ST45<br>genomes | 100.00 | 100.00 | Lorraine FQ958210.1 |  |
| 300251<br>1 | 300415<br>4 | 1644 | All ST45<br>genomes | 100.00 | 100.00 | Lorraine FQ958210.1 |  |
| 302288<br>9 | 302511<br>3 | 2225 | All ST45<br>genomes | 100.00 | 100.00 | PATHC002<br>CP113439.1 | Homology to<br>ISLpn9 |
| 313594<br>3 | 313718<br>5 | 1243 | All ST45<br>genomes | 100.00 | 100.00 | Lorraine FQ958210.1 |  |
| 321092<br>7 | 321209<br>0 | 1164 | Very low<br>coverage in<br>Phildelphia<br>reference | 100.00 | 98.63 | Lorraine FQ958210.1 | Homology to<br>ISLpn9 |
| 332200<br>5 | 332300<br>6 | 1002 | Very low<br>coverage in<br>Phildelphia<br>reference | 100.00 | 99.90 | Lorraine FQ958210.1 | Homology to<br>ISLpn9 |
| 338047<br>4 | 338147<br>4 | 1001 | Very low<br>coverage in<br>Phildelphia<br>reference | 100.00 | 99.70 | Lorraine FQ958210.1 | Homology to<br>ISLpn9 |
| 339557<br>1 | 339844<br>6 | 2876 | All ST45<br>genomes | 100.00 | 100.00 | Lorraine FQ958210.1 |  |
| 340547<br>3 | 340663<br>4 | 1162 | All ST45<br>genomes | 100.00 | 100.00 | Lorraine FQ958210.1 |  |
| 341170<br>0 | 341303<br>8 | 1339 | All ST45<br>genomes | 100.00 | 100.00 | Lorraine FQ958210.1 |  |
| 341349<br>0 | 341674<br>6 | 3257 | All ST45<br>genomes | 100.00 | 100.00 | Lorraine FQ958210.1 |  |
| 346290<br>8 | 346422<br>9 | 1322 | All ST45<br>genomes | 100.00 | 99.62 | Lorraine FQ958210.1 | Homology to<br>ISLpn9 |
| 349427<br>7 | 349590<br>4 | 1628 | Very low<br>coverage in<br>Phildelphia<br>reference | 100.00 | 99.63 | PATHC002<br>CP113439.1 | Homology to<br>ISLpn9 |
| 352048<br>9 | 352192<br>1 | 1433 | All ST45<br>genomes | 94.00 | 99.93 | H3 P114576.1 |  |

## References

1. Fields BS, Benson RF, Besser RE. *Legionella* and Legionnaires’ Disease: 25 Years of Investigation. Clin Microbiol Rev. 2002 Jul;15(3):506–26.

2. Gao LY, Harb OS, Abu Kwaik Y. Utilization of similar mechanisms by Legionella pneumophila to parasitize two evolutionarily distant host cells, mammalian macrophages and protozoa. Infect Immun. 1997 Nov;65(11):4738–46.

3. Chauhan D, Shames SR. Pathogenicity and Virulence of *Legionella*: Intracellular replication and host response. Virulence. 2021 Dec 31;12(1):1122–44.

4. Wuthrich D, Gautsch S, Spieler-Denz R, Dubuis O, Gaia V, Moran-Gilad J, et al. Air-conditioner cooling towers as complex reservoirs and continuous source of *Legionella pneumophila* infection evidenced by a genomic analysis study in 2017, Switzerland. Euro surveillance: bulletin Europeen sur les maladies transmissibles = European communicable disease bulletin. 2019 Jan;24(4).

5. Hilbi H, Buchrieser C. Microbe Profile: Legionella pneumophila - a copycat eukaryote: This article is part of the Microbe Profiles collection. Microbiology [Internet]. 2022 Mar 1 [cited 2024 Oct 21];168(3). Available from: https://www.microbiologyresearch.org/content/journal/micro/10.1099/mic.0.001142

6. Molmeret M, Horn M, Wagner M, Santic M, Abu Kwaik Y. Amoebae as Training Grounds for Intracellular Bacterial Pathogens. Appl Environ Microbiol. 2005 Jan;71(1):20–8.

7. Van Kenhove E, Dinne K, Janssens A, Laverge J. Overview and comparison of Legionella regulations worldwide. American Journal of Infection Control. 2019 Aug;47(8):968–78.

8. Bundesamt für Lebensmittelsicherheit und Veterinärwesen. Empfehlungen zu Legionellen und Legionellose [Internet]. 2024 [cited 2024 Aug 13]. Available from: https://www.blv.admin.ch/blv/de/home/gebrauchsgegenstaende/badewasser/empfehlungen-legionellen-legionellose.html

9. Dahlen G. Biofilms in Dental Unit Water Lines. In: Eick S, editor. Monographs in Oral Science [Internet]. S. Karger AG; 2021 [cited 2024 Jul 26]. p. 12–8. Available from: https://karger.com/books/book/383/chapter/5565067

10. Petti S, Vitali M. Occupational risk for *Legionella* infection among dental healthcare workers: meta-analysis in occupational epidemiology. BMJ Open. 2017 Jul;7(7):e015374.

11. Bartley PB, Ben Zakour NL, Stanton-Cook M, Muguli R, Prado L, Garnys V, et al. Hospital-wide Eradication of a Nosocomial *Legionella pneumophila* Serogroup 1 Outbreak. Clin Infect Dis. 2016 Feb 1;62(3):273–9.

12. Rangel-Frausto MS, Rhomberg P, Hollis RJ, Pfaller MA, Wenzel RP, Helms CM, et al. Persistence of Legionella pneumophila in a hospital’s water system: a 13-year survey. Infect Control Hosp Epidemiol. 1999 Dec;20(12):793–7.

13. Perola O, Kauppinen J, Kusnetsov J, Kärkkäinen UM, Lück PC, Katila ML. Persistent Legionella pneumophila colonization of a hospital water supply: efficacy of control methods and a molecular epidemiological analysis. APMIS. 2005 Jan;113(1):45–53.

14. Croucher NJ, Page AJ, Connor TR, Delaney AJ, Keane JA, Bentley SD, et al. Rapid phylogenetic analysis of large samples of recombinant bacterial whole genome sequences using Gubbins. Nucleic Acids Research. 2015;43(3):e13.

15. Hadfield J, Croucher NJ, Goater RJ, Abudahab K, Aanensen DM, Harris SR. Phandango: an interactive viewer for bacterial population genomics. Bioinformatics. 2017;34(2):292–3.

16. Didelot X, Croucher NJ, Bentley SD, Harris SR, Wilson DJ. Bayesian inference of ancestral dates on bacterial phylogenetic trees. Nucleic Acids Res. 2018 Dec 14;46(22):e134.

17. Feldgarden M, Brover V, Haft DH, Prasad AB, Slotta DJ, Tolstoy I, et al. Validating the AMRFinder Tool and Resistance Gene Database by Using Antimicrobial Resistance Genotype-Phenotype Correlations in a Collection of Isolates. Antimicrob Agents Chemother. 2019 Nov;63(11).

18. Wüthrich, Daniel, Gautsch S, Spieler-Denz R, Dubuis O, Gaia V, Moran-Gilad J, et al. Air-conditioner cooling towers as complex reservoirs and continuous source of Legionella pneumophila infection evidenced by a genomic analysis study in 2017, Switzerland. Eurosurveillance. 2019 Jan 24;24(4):1800192.

19. Haviernik J, Dawson K, Anderson T, Murdoch D, Chambers S, Biggs P, et al. Complete Genome Sequence of a Legionella longbeachae Serogroup 2 Isolate Derived from a Patient with Legionnaires’ Disease. Putonti C, editor. Microbiol Resour Announc. 2020 Jan 30;9(5):10.1128/mra.01563-19.

20. Bédard E, Paranjape K, Lalancette C, Villion M, Quach C, Laferrière C, et al. Legionella pneumophila levels and sequence-type distribution in hospital hot water samples from faucets to connecting pipes. Water Research. 2019 Jun;156:277–86.

21. Lepine LA, Jernigan DB, Butler JC, Pruckler JM, Benson RF, Kim G, et al. A recurrent outbreak of nosocomial legionnaires’ disease detected by urinary antigen testing: evidence for long-term colonization of a hospital plumbing system. Infect Control Hosp Epidemiol. 1998 Dec;19(12):905–10.

22. Byrne BG, McColm S, McElmurry SP, Kilgore PE, Sobeck J, Sadler R, et al. Prevalence of Infection-Competent Serogroup 6 *Legionella pneumophila* within Premise Plumbing in Southeast Michigan. Shuman HA, editor. mBio. 2018 Mar 7;9(1):e00016–18.

23. Casini B, Valentini P, Baggiani A, Torracca F, Frateschi S, Nelli LC, et al. Molecular epidemiology of Legionella pneumophila serogroup 1 isolates following long-term chlorine dioxide treatment in a university hospital water system. Journal of Hospital Infection. 2008 Jun;69(2):141–7.

24. Oberdorfer K, Müssigbrodt G, Wendt C. Genetic diversity of Legionella pneumophila in hospital water systems. International Journal of Hygiene and Environmental Health. 2008 Mar;211(1–2):172–8.

25. Pancer K, Matuszewska R, Bartosik M, Kacperski K, Krogulska B. Persistent colonization of 2 hospital water supplies by L. pneumophila strains through 7 years--sequence-based typing and serotyping as useful tools for a complex risk analysis. Ann Agric Environ Med. 2013;20(4):687–94.

26. Gorzynski J, Wee B, Llano M, Alves J, Cameron R, McMenamin J, et al. Epidemiological analysis of Legionnaires’ disease in Scotland: a genomic study. The Lancet Microbe. 2022 Nov;3(11):e835–45.

27. David S, Afshar B, Mentasti M, Ginevra C, Podglajen I, Harris SR, et al. Seeding and Establishment of Legionella pneumophila in Hospitals: Implications for Genomic Investigations of Nosocomial Legionnaires’ Disease. Clinical Infectious Diseases. 2017 May 1;64(9):1251–9.

28. David S, Mentasti M, Lai S, Vaghji L, Ready D, Chalker VJ, et al. Spatial structuring of a Legionella pneumophila population within the water system of a large occupational building. Microbial Genomics [Internet]. 2018 Oct 1 [cited 2024 Apr 19];4(10). Available from: https://www.microbiologyresearch.org/content/journal/mgen/10.1099/mgen.0.000226

29. Geschichte der Zahnmedizin an der Universität Basel seit 1888 [Internet]. Universität Basel; [cited 2024 Sep 6]. Available from: https://unigeschichte.unibas.ch/fakultaeten-und-faecher/medizinische-fakultaet/juengste-entwicklungen-der-medizinischen-fakultaet/zahnmedizin

30. Orsi GB, Vitali M, Marinelli L, Ciorba V, Tufi D, Del Cimmuto A, et al. Legionella control in the water system of antiquated hospital buildings by shock and continuous hyperchlorination: 5 years experience. BMC Infect Dis. 2014 Dec;14(1):394.

31. Lombardi A, Borriello T, De Rosa E, Di Duca F, Sorrentino M, Torre I, et al. Environmental Monitoring of Legionella in Hospitals in the Campania Region: A 5-Year Study. IJERPH. 2023 Apr 14;20(8):5526.

32. Lee HK, Shim JI, Kim HE, Yu JY, Kang YH. Distribution of *Legionella* Species from Environmental Water Sources of Public Facilities and Genetic Diversity of *L. pneumophila* Serogroup 1 in South Korea. Appl Environ Microbiol. 2010 Oct;76(19):6547–54.

33. Tijet N, Tang P, Romilowych M, Duncan C, Ng V, Fisman DN, et al. New Endemic *Legionella pneumophila* Serogroup I Clones, Ontario, Canada. Emerg Infect Dis. 2010 Mar;16(3):447–54.

34. Qin T, Zhou H, Ren H, Guan H, Li M, Zhu B, et al. Distribution of Sequence-Based Types of Legionella pneumophila Serogroup 1 Strains Isolated from Cooling Towers, Hot Springs, and Potable Water Systems in China. Elkins CA, editor. Appl Environ Microbiol. 2014 Apr;80(7):2150–7.

35. Zhan XY, Zhu QY. Molecular typing of Legionella pneumophila isolates from environmental water samples and clinical samples using a five-gene sequence typing and standard Sequence-Based Typing. Bevivino A, editor. PLoS ONE. 2018 Feb 1;13(2):e0190986.

36. Echaihidi F, Prevost B, Martiny D, Wybo I, Piérard D, Michel C. Activity report from 2011 to 2022 Reference centre for Legionella pneumophila UZ Brussel – LHUB-ULB [Internet]. National Reference Centre for Legionella pneumophila, Belgium; 2022 [cited 2024 Aug 26]. Available from: https://www.sciensano.be/sites/default/files/legionella_2011-2022_nrc_rapport_english_final.pdf

37. Monjo MP. Structure and virulence of Legionella pneumophila populations from freshwater systems in Germany and Middle East [Internet]. Technischen Universität Carolo-Wilhelmina zu Braunschweig; 2016 [cited 2024 Aug 26]. Available from: https://leopard.tu-braunschweig.de/servlets/MCRFileNodeServlet/dbbs_derivate_00043159/Diss_Pecellin_Marina.pdf

38. Fotos PG, Westfall HN, Snyder IS, Miller RW, Mutchler BM. Prevalence of Legionella-specific IgG and IgM antibody in a dental clinic population. J Dent Res. 1985 Dec;64(12):1382–5.

39. Gomez-Valero L, Rusniok C, Jarraud S, Vacherie B, Rouy Z, Barbe V, et al. Extensive recombination events and horizontal gene transfer shaped the Legionella pneumophila genomes. BMC Genomics. 2011 Dec;12(1):536.

40. David S, Sánchez-Busó L, Harris SR, Marttinen P, Rusniok C, Buchrieser C, et al. Dynamics and impact of homologous recombination on the evolution of Legionella pneumophila. Didelot X, editor. PLoS Genet. 2017 Jun 26;13(6):e1006855.

